# Cell wall strengthening by phenylpropanoid dehydrodimers during the plant hypersensitive cell death

**DOI:** 10.1101/2022.12.20.521293

**Authors:** Basem Kanawati, Marko Bertic, Franco Moritz, Felix Habermann, Ina Zimmer, David Mackey, Philippe Schmitt-Kopplin, Jörg-Peter Schnitzler, Jörg Durner, Frank Gaupels

**Author notes:** These authors contributed equally to this article (Basem Kanawati, Marko Bertic). Author for communication: Dr. Frank Gaupels.

## Abstract

Infection of Arabidopsis with avirulent *Pseudomonas syringae* and exposure to nitrogen dioxide (NO_2_) both trigger hypersensitive cell death (HCD) that is characterized by the emission of bright blue-green (BG) autofluorescence under UV illumination. The aim of our current work was to identify the BG fluorescent molecules and scrutinize their biosynthesis and functions during the HCD. Compared to wild-type (WT) plants, the phenylpropanoid-deficient mutant *fah1* developed normal HCD except for the absence of BG fluorescence. Ultrahigh resolution metabolomics combined with mass difference network analysis revealed that WT but not *fah1* plants rapidly accumulate dehydrodimers of sinapic acid, sinapoylmalate, 5-OH-ferulic acid, and 5-OH-feruloylmalate during the HCD. FAH1-dependent BG fluorescence appeared exclusively within dying cells of the upper epidermis as detected by microscopy. Saponification released dehydrodimers from extracted cell wall material. Collectively, our data suggest that HCD induction leads to the formation of free BG fluorescent dehydrodimers from monomeric sinapates and 5-hydroxyferulates. Reactive oxygen species from de-regulated photosynthesis likely contribute to the radical-radical coupling. The formed dehydrodimers move from upper epidermis cells into the apoplast where they esterify and thereby cross-link cell wall polymers. Both, free as well as wall-bound phenylpropanoid dehydrodimers are defense-related compounds in Arabidopsis. We propose that other plants also employ dehydrodimers of highly abundant phenylpropanoids for rapid defense against pathogen attack.

## INTRODUCTION

Infection of Arabidopsis thaliana with avirulent *Pseudomonas syringae* pv. *tomato* expressing the avirulence gene *AvrRpm1* (*P*.*s*.*t. AvrRpm1*) triggers the hypersensitive defence response (HR) culminating in programmed cell death (PCD) (Katagiri et al., 2002; Coll et al., 2011). HR-PCD can be distinguished from other forms of PCD by the appearance of UV-induced blue-green (BG) fluorescence in dying leaf tissues (Nicholson and Hammerschmidt, 1992; Bennett et al., 1996). HR-PCD depends on simultaneous signaling by reactive oxygen- and nitrogen species (ROS and RNS) (Delledonne et al., 2001; Gaupels et al., 2011). Exposure of Arabidopsis to the RNS nitrogen dioxide (NO_2_) results in basal pathogen immunity or rapid cell death in a dose-dependent manner (Kasten et al., 2016; Mayer et al., 2018). NO_2_-induced cell death is accompanied by the strong emission of BG fluorescence under UV light similar to HR-PCD (Frank et al., 2019). Hereafter, NO_2_- and *P*.*s*.*t. AvrRpm1*-induced cell death will be collectively called hypersensitive cell death (HCD).

The enigmatic BG fluorescence under UV illumination was recognized long ago (Nicholson and Hammerschmidt, 1992). However, even after more than 40 years of intense research the source of the fluorescence remained ambiguous. Based on microscopy and spectral analyses phenolic compounds such as phenylpropanoids might be a major source of autofluorescence during the HCD without individual molecules identified to date (Nicholson and Hammerschmidt, 1992). In Arabidopsis sinapic acid (Si) and sinapoylmalate (SM) are the dominant phenylpropanoids mainly localized in the leaf upper epidermis where they act as a sunscreen by scavenging harmful UV irradiation under emission of blue fluorescence (Chapple et al., 1992; Milkowski and Strack, 2010).

These findings prompted us to investigate HCD in mutants deficient in phenylpropanoids. The results of our study suggest that derivatives of Si and its precursor 5-hydroxyferulic acid (Hf) form BG fluorescent dehydrodimers. Additionally, we scrutinized the mechanism of formation and possible functions of the phenylpropanoid dehydrodimers.

## RESULTS

### BG autofluorescence is dependent on sinapates and 5-hydroxyferulates

Leaves of untreated Arabidopsis plants emit red chlorophyll fluorescence and blue fluorescence under UV illumination (Figure 1A). The blue fluorescence originates from sinapates localized in the upper leaf epidermis (Milkowski and Strack, 2010). For the investigation of HR-PCD, we used transgenic Arabidopsis expressing the *P*.*s*.*t*. avirulence gene *AvrRpm1* under control of a dexamethasone (Dex)-inducible promoter which allows the reproducible initiation of HR without a pathogen (Mackey et al., 2002). Spraying *Dex::AvrRpm1* plants with Dex elicited rapid cell death, leaf collapse, and the emission of bright BG fluorescence at 3 h after treatment (Figure 1B). Fumigation for 1 h with 30 ppm NO_2_ led to rapid cell death, leaf collapse and strong BG fluorescence immediately after the treatment (Figure, 1C and D).

**Figure 1.**
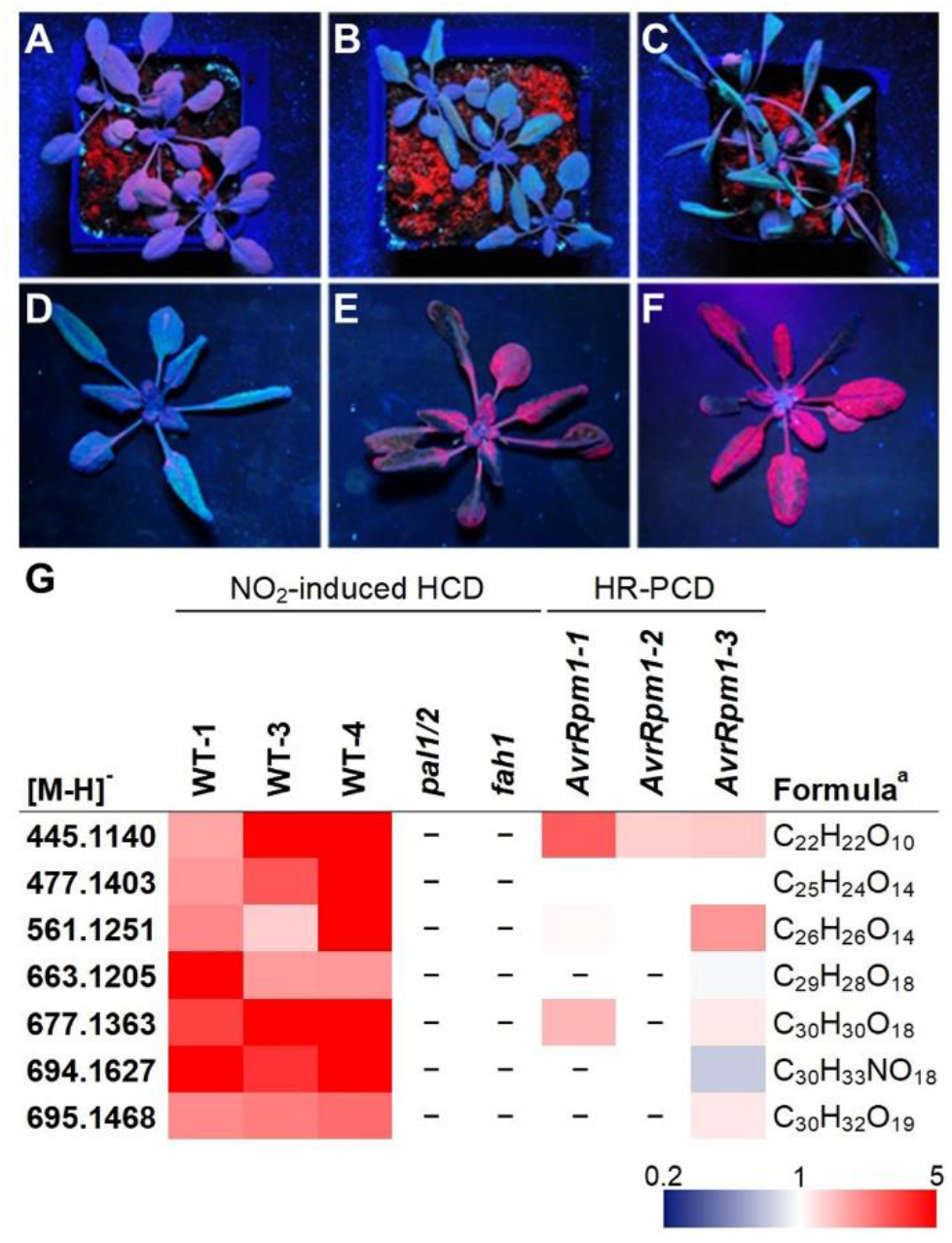
The mutants *fah1* and *pal1/2* do not show BG fluorescence during the HCD. Plants were observed under UV light for fluorescence emission from chlorophyll (red) and phenolic secondary metabolites (BG). A, Untreated Arabidopsis wild-type (WT) plants. B, *Dex::AvrRpm1* transgenic plants 3 h after induction of HR-PCD by spraying dexamethasone solution. C, and (D) Arabidopsis WT plants immediately after fumigation for 1 h with 30 ppm NO_2_. E, *pal1/2* and (F) *fah1* mutant plants after NO_2_ treatment. *pal1/2* has reduced levels of phenylpropanoids. *fah1* is deficient in Hf and Si. G, Leaf extracts were analyzed by non-targeted FT-ICR-MS. The figure depicts unknown metabolites accumulating in all three NO_2_ fumigation experiments but not detected *pal1/2* or *fah1*. Red and blue shades indicate significantly increased or decreased spectral counts (fold change), respectively. White color indicates not significantly regulated and white plus dash not detected. Please refer to Suppelmental File S1 for original data including statistics. ^a^, sum formula of uncharged molecule.

By contrast, the knock-out mutants *pal1/2* (Figure 1E) and *fah1* (Figure 1F) undergo NO_2_-induced PCD but do not show blue or BG fluorescence when inspected under UV. *pal1/2* has low levels of phenylpropanoids due to mutation of *PAL1* and *PAL2* (Rohde et al., 2004). Both genes code for phenylalanine ammonia lyases (PAL) that catalyze the initial reaction of the phenylpropanoid pathway, namely the synthesis of cinnamic acid from phenylalanine (Fraser and Chapple, 2011). The PAL activity is reduced by 75% in the *pal1/2* double mutant (Rohde et al., 2004). *fah1* is deficient in Hf and sinapates due to mutation of the gene *FAH1* coding for the enzyme ferulate-5-hydroxylase that converts ferulic acid into Hf, the latter being subsequently O-methylated to Si (Fraser and Chapple, 2011). Hence, BG fluorescence in dying leaf tissues is dependent on Hf and Si. However, the induction of HCD is independent of BG fluorescence, Hf, and Si.

### Radical-mediated dehydrodimerization of phenylpropanoids is a source of BG fluorescence

The *pal1/2* and *fah1* mutant lines were further employed to identify BG fluorescent metabolites by direct-infusion Fourier transform ion cyclotron resonance mass spectrometry (FT-ICR-MS), a technique that facilitates the determination of molecular masses with sub-ppm accuracy (<0.001 Da). Accurate masses can then be used to calculate the chemical formulae and generate putative annotations. This non-targeted metabolomics approach detected seven metabolites that showed increased levels after NO_2_ exposures of WT plants but were absent in *fah1* and *pal1/2* (Figure 1G). One metabolite with the negative ion mass ([M-H]^-^) 445.1140 Da accumulated also upon induction of HR-PCD. In previous studies compounds with similar measured masses were identified as sinapate dehydrodimers (Bunzel et al., 2003; Frolov et al., 2013; Yin et al., 2019) suggesting that HCD is accompanied by dimer formation between phenylpropanoids and, more specifically, between 5-hydroxyferulates and sinapates. SM, Si, 5-hydroxyferuloylmalate (HM), and Hf are the most abundant FAH1-dependent metabolites in Arabidopsis leaf extracts (Supplemental File S1).

Dehydrodimerization involves peroxidase-mediated one electron oxidation of the monomeric precursors to give dehydrogenated radicals (Ralph et al., 2004; Bunzel, 2010). Therefore, the FT-ICR-MS data were further scrutinized by mass difference network (MDN) analysis assuming that a dehydroradical of either SM, Si, HM, or Hf reacted with another unknown radical (Figure 2A). This approach uncovered possible molecular interactions (Supplemental File S1). Three generated networks contained metabolites that were consistently regulated in all NO_2_ fumigation experiments (Figure 2A). According to network I the compound with the sum formula C_30_H_32_O_19_ could arise from dehydrodimerization of Si, SM, or HM radicals with three different binding partners. C_16_H_20_O_10_ might represent 5-hydroxyferulate glucose (HG) since it does not occur in *fah1*. In this case, C_30_H_32_O_19_ that is also absent in *fah1* could refer to the HM-HG dehydrodimer. In network II both, the precursor C_15_H_19_NO_9_ as well as the putative dehydrodimer C_30_H_33_NO_18_ did not result in meaningful database hits.

**Figure 2.**
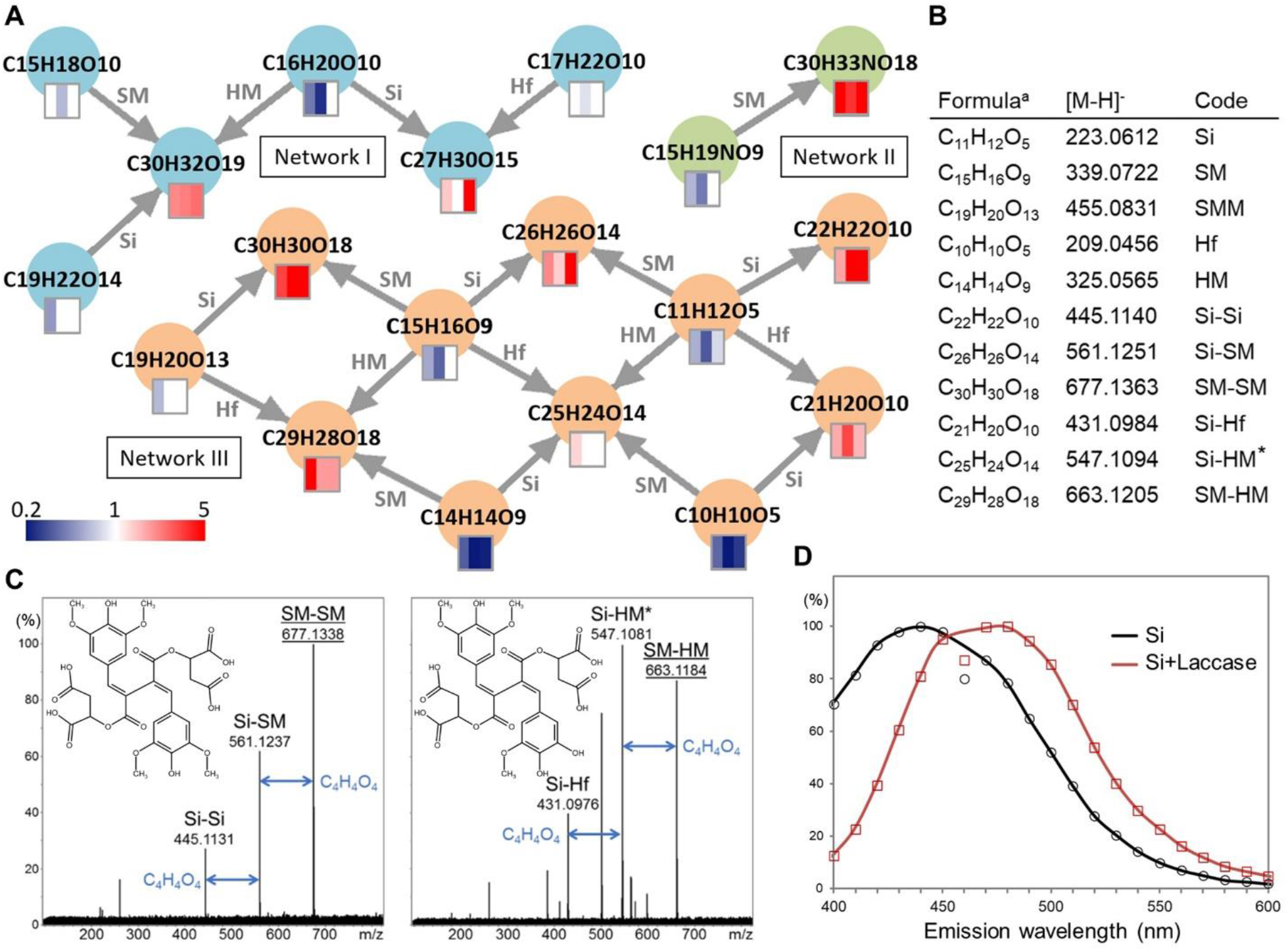
Dehydrodimers of sinapates and 5-hydroxyferulates accumulate during NO_2_-induced cell death. A, Mass difference network (MDN) analysis. Accurate negative ion masses as determined by FT-ICR-MS were analyzed for mass differences due to binding of dehydrogenated Si, SM, Hf, and HM (for complete datasets see Supplemental File S1). The node colors indicate different sub-networks. Square icons depict the NO_2_-induced regulation in three independent experiments. The color code represents fold change in spectral count (NO_2_/Air ratio). White color means not detected or not regulated (for data and statistics see Supplemental File S1). B, Chemical formulae of the uncharged molecules, measured accurate masses of the [M-H]^-^ ions, and code describing the monomers and dimers found in network III shown in (A). C, MS/MS fragmentation patterns at a collision energy of 8 eV of the SM-SM and SM-HM dehydrodimers. Parent ions are underlined. Representative structures of 8-8-coupled SM-SM and SM-HM dimers are shown. The loss of malyl (C_4_H_4_O_4_) moieties during the fragmentation is indicated by blue arrows. The full peak lists can be found in Figure S1. D, Fluorescence spectra of Si alone and Si after laccase addition causing the formation of Si-Si dehydrodimers. The excitation wavelength was 320 nm. *, Si-HM or SM-Hf. ^a^, sum formula of uncharged molecule.

The monomeric precursors depicted in network III likely correspond to Si, SM, sinapoyldimalate (SMM), Hf, and HM (Figure, 2A and B) because they were highly abundant in WT but absent or strongly depleted in *fah1* and *pal1/2* (Supplemental File S1). The spectral counts of the monomers mostly decreased upon NO_2_ fumigation. MS/MS fragmentation analyses confirmed the identities of SM and SMM (Supplemental Figure S1, A and B). Network III suggests the formation of Si-Si, Si-SM, SM-SM, Si-Hf, Si-HM or SM-Hf (isomers not distinguishable), and SM-HM upon HCD induction (Figure, 2A and B). Five dehydrodimers accumulated up to 10-fold in all experiments whereas the levels of Si-HM/SM-Hf were generally low (Supplemental File S1). MS/MS fragmentation analysis confirmed the identity of the SM-SM and SM-HM dimers (Figure 2C, Supplemental Figure S1, C and D), the most prominent fragmentation pattern being losses of the malyl moieties. Dehydrodimers of ferulates were not regulated during the HCD (Supplemental File S1).

Next, we assessed whether the rise of dehydrodimers could explain the observed BG fluorescence emission during HCD. Peroxidases and laccase are known to catalyze polymerisations of phenolic compounds (Ralph et al., 2004; Tufegdžić et al., 2005). Si fluoresced blue under UV-A (320 nm) illumination. However, upon addition of laccase the fluorescence turned BG (Figure 2D). The pronounced shift of the fluorescence maximum from 440 nm to 470 nm probably reflects dimerization of Si and is likely due to the increased size of the π conjugated system in the formed dehydrodimers. A similar shift in fluorescence was previously observed upon dehydrodimerization of ferulate (Tufegdžić et al., 2005).

### Fluorescent phenylpropanoid dehydrodimers arise within epidermal cells next to dying mesophyll cells

Untreated plants displayed UV-excited blue fluorescence in the adaxial epidermis but not the abaxial epidermis that lacks Si and Hf derivatives (Supplemental Figure S2, A and B). Fumigation with 30 ppm NO_2_ for 20 min led to a transient increase in red fluorescence due to disturbed photosynthesis (Mayer et al., 2018) and rapid development of BG fluorescent lesions (Figure 3A). At 24 h after treatment cells at the edge of the lesion appeared bright blue whereas cells in the center of the lesion were BG fluorescent or non-fluorescent (Figure, 3A and B). Mesophyll cells turned chlorotic during the HCD (Figure 3C). The bright blue fluorescence was particularly high in the cell peripheries (Figure 3D). Close examinations revealed that the BG fluorescence in the lesion centers vanished during HCD progression (Figure 3E). Infection with *P*.*s*.*t. AvrRpm1* also triggered the emission of bright blue and BG fluorescence during the HR (Figure 3F). Both, the bright blue as well as BG fluorescence were not observed in *fah1* (Figure, 3G-I) suggesting that they are related to the dehydrodimerization of Si- and Hf derivatives (Figures 1 and 2). It is yet unclear whether the BG and bright blue fluorescence emissions are related to different FAH1-dependent dehydrodimers or other factors such as chemical modifications, cellular localization or optical effects.

**Figure 3.**
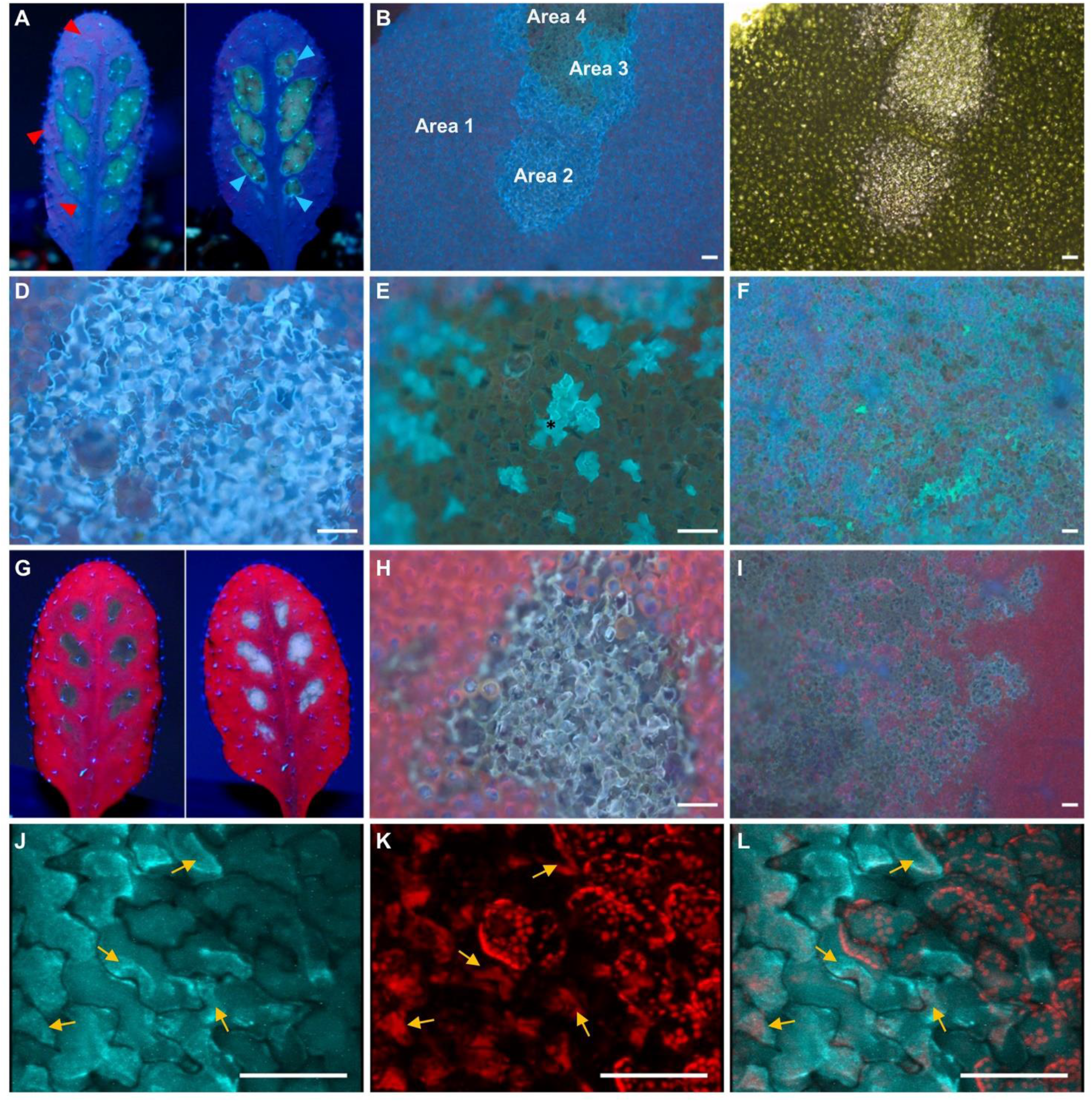
HCD-related BG and bright blue fluorescence is mainly localized in upper epidermal cells next to dying mesophyll cells. UV-induced fluorescence imaging by (A and G) a handheld UV lamp plus camera, (B-F, H and I) fluorescence microscope, and (J-L) CLSM. HCD was triggered by fumigation with 30 ppm NO_2_ for 20 min or infection with *P*.*s*.*t. AvrRpm1*. A, Leaf of a WT plant at 4 h (left panel) and 24 h (right panel) after NO_2_ exposure. Note the transiently enhanced red chlorophyll fluorescence at 4 h (red arrow heads) and the bright blue fluorescence (blue arrow heads) surrounding the lesions at 24 h after treatment. B, Fluorescence emission under UV-A (365 nm) excitation. Weak blue fluorescence was visible in unaffected tissues (area 1). Within the lesion epidermal cells fluoresced bright blue (area 2) or BG (area 3). Dead tissue appeared brown (area 4). C, Transmission light revealed the chlorosis of mesophyll cells within the lesion. D, HCD-related bright blue fluorescence was strongest in the peripheries of epidermal cells. E, During HCD progression fluorescence in epidermal cells turned BG and subsequently vanished. For instance, the cell highlighted by an asterisk was already partially transparent. Dead epidermis cells were transparent but mesophyll cells brown probably due to chlorophyll degradation products. F, *P*.*s*.*t. AvrRpm1*-induced fluorescence in WT at 24 h after infection. BG and bright blue fluorescence was absent in *fah1* undergoing (G and H) NO_2_-induced or (I) *P*.*s*.*t. AvrRpm1*-induced HCD. (J-L) Tissue at the lesion edges were examined by CLSM. J, Localization of bright blue fluorescence (excitation at 364 nm/emission at ≥385 nm) and (K) red chlorophyll fluorescence (exc. at 633 nm/emi. at ≥650 nm), and (L) digital overlay of both (arbitrary colors). (J-L) Bright blue fluorescence often appeared at contact sites (arrows) of epidermis cells with mesophyll cells undergoing PCD as evidenced by broken chloroplasts. However, the bright blue fluorescence did not turn up in mesophyll cells or in epidermis cells next to live mesophyll cells with intact chloroplasts (refer also to Movies 1 and 2). Scale bars indicate 100 µm.

Bathing leaf tissues in a highly osmotic solution uncovered that the BG fluorescence was localized within the shrinking protoplast (Supplemental Figure S2, C and D). Hence, dehydrodimers mainly arise in the vacuoles or the cytoplasm. We further analyzed in more detail the bright blue fluorescence at the lesion edges by confocal laser scanning microscopy (CLSM) which revealed a major source of UV-excited fluorescence at contact sites between epidermis cells and dying mesophyll cells (Figure, 3J-L, Movies 1 and 2). Burst chloroplasts in the mesophyll always co-localized with bright blue fluorescence.

### Phenypropanoid dehydrodimers function in inducible cell wall cross-linking during the HCD

It was shown before that dehydrodimers are bound to the cell wall by esterification. To test wether this also occurs in Arabidopsis we saponified cell wall fractions and analyzed the extracted esters by ultra performance liquid chromatograpy-tandem MS (UPLC-MS/MS). HCD induction by NO_2_ or *P*.*s*.*t. AvrRpm1* caused the attachment of SM-SM, SM-Si, and Si-Si to the cell wall via ester bonds whereas these dimers were not detected in leaf extracts from control plants (Figure 4A). Ferulate appeared to be a constitutive cell wall ester not showing up-regulated levels upon HCD induction (Figure 4A). SM-SM esterified completely and Si-Si as well as Si-SM partially to cell wall polymers wheres SM-Hf/Si-HM accumulated exclusively in its free form until 24 h after NO_2_ treatment (Figure 4B).

**Figure 4.**
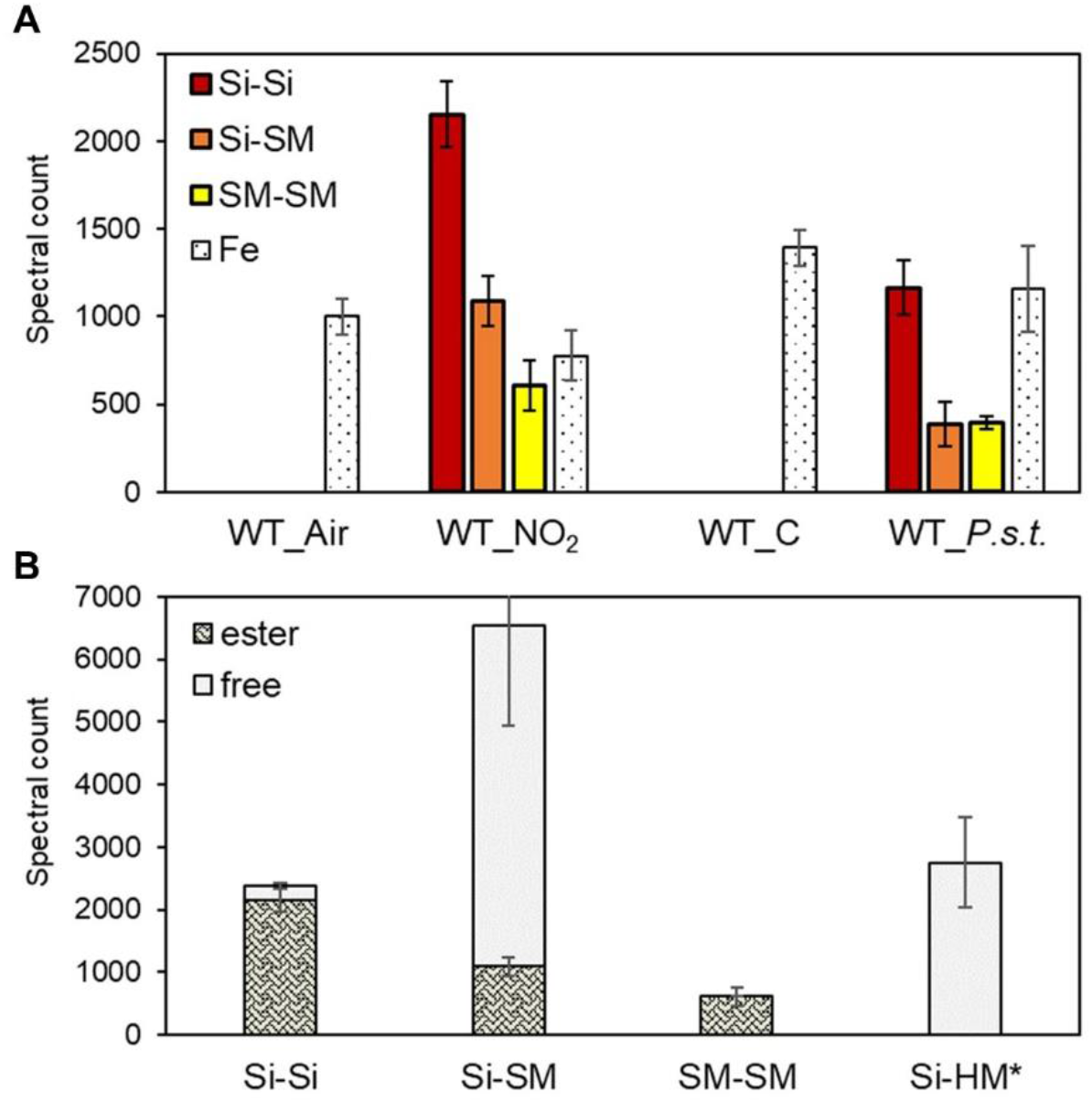
Dehydrodimers partially esterify to the cell wall during the HCD. A, The dehydrodimers and ferulate (Fe) were released from purified cell wall material by saponification before analysis by UPLC-MS/MS. Semi-quantification of dehydrodimer esters in wild-type (WT) plants 24 h after fumigation with NO_2_ or infection with avirulent *P*.*s*.*t. AvrRpm1* (*P*.*s*.*t*.). B, Some dehydrodimers only partially or not at all esterified to the cell wall within 24 h after NO_2_ fumigation. Error bars depict standard deviations (n = 6).

## DISCUSSION

Dehydrodimers of sinapates and 5-hydroxyferulates are sources of the enigmatic BG fluorescence emerging during the HCD in Arabidopsis, which is supported by three observations: (1.) The Si- and Hf-deficient mutant *fah1* does not exhibit BG fluorescence. (2.) Dehydrodimers are the only metabolites that accumulate during the HCD in a FAH1-dependent manner. (3.) Laccase-mediated oligomerization of Si *in vitro* causes a shift from blue to BG fluorescence as it also occurs in dying epidermis cells.

Ferulates were previously found to be the major but sinapates only minor precursors of dehydrodimers in various investigated plant species (Buanafina, 2009; Santiago and Malvar, 2010). Dehydrodiferulates facilitate cell wall cross-linking during growth and development. Available evidence additionally argues for a role of these molecules in plant defence responses. For example, in different maize cultivars the levels of dehydrodiferulates exhibited a positive correlation with resistance against the fungal pathogen *Fusarium graminearum* (Bily et al., 2003). Ikegawa et al. (Ikegawa et al., 1996) observed an increased esterification of hemicellulose by dehydrodiferulates during the HR of oat (*Avena sativa*) towards *Puccinia coronata* f.sp. *avenae*. And transgenic *Brachypodium distachyon* expressing a fungal ferulic acid esterase (FAE) had reduced levels of ferulate and diferulate esters corresponding with an enhanced susceptibility to *Botrytis cinerea* (Reem et al., 2016).

Our current research provides new insights in the inducible formation of phenylpropanoid dehydrodimers during plant defence responses. HCD induction did not cause the dimerization of ferulates in Arabidopsis. Instead of that UV protectant sinapates and 5-hydroxyferulates were repurposed to promote rapid cell wall cross-linking in the upper epidermis probably as an effective defence against air-born pathogens. Future research will reveal whether highly abundant phenypropanoids in other plant species such as syringaldehyde and chlorogenic acid in lettuce (Bennett et al., 1996) dimerize during the HCD.

It is generally assumed that – at least in monocots – ferulate monomers esterify to polysaccharides such as pectin already within the cell serving as building blocks for cell wall fortification (Buanafina, 2009). However, according to our recent research neither monomers nor dimers of sinapates and 5-hydroxyferulates are attached to polysaccharides in Arabidopsis. The exact localization of dehydrodiferulate formation is still debated. Dimers could form intracellularly, extracellularly and/or *in muro* (Lindsay and Fry, 2008; Buanafina, 2009). We found a strong intracellular source of BG fluorescence arguing for dimerization of sinapates and 5-hydroxyferulates within the upper epidermis cells. In addition to the prominent BG fluorescence a FAH1-dependent bright blue fluorescence appeared at the lesion edges. CLSM uncovered that burst chloroplasts in dying mesophyll cells always co-localized with spots of bright blue-fluorescence in nearby epidermis cells. Thus, ROS accumulation caused by disrupted photosynthesis could contribute to the formation of dehydrodimers at contact sites between dying mesophyll- and epidermis cells.

Individual dehydrodimers exhibited markedly different wall-binding kinetics. Si-Si and SM-SM mostly esterified to the cell wall whereas the pools of Si-SM remained largely and Si-HM/SM-Hf completely unbound until 24 h after onset of the HCD. These findings point to specific functions of free versus wall-bound dehydrodimers in pathogen defence. We did not detect diferulate in Arabidopis. Ferulate monomers were not regulated during the HCD indicating that they might contribute to cell wall cross-linking during growth and development rather than defence responses.

In conclusion, HCD induction leads to the rapid formation of BG fluorescent dehydrodimers from large pre-existing pools of UV protectant Si and Hf derivatives within dying epidermis cells. The dimerization could be mediated by ROS from burst chloroplasts in the mesophyll. At least a proportion of the dimers Si-Si, Si-SM, SM-SM move into the apoplast where they bind to cell wall polymers by esterification thereby turning non-fluorescent. Dehydrodimer formation is not essential for the execution of HCD but is rather related to cell wall reinforcement. Additionally, free Si-HM/SM-Hf and other phenylpropanoid dehydrodimers could have so far unknown functions in pathogen defence.

## MATERIALS AND METHODS

### Plant material and HCD induction

*Arabidopsis thaliana* ecotype Columbia-0 (Col-0), transgenic *pDex::AvrRpm1-HA* (Mackey et al., 2002), *fah1* (Chapple et al., 1992), and *pal1 pal2* (Rohde et al., 2004) plants were grown for 4-5 weeks on soil under long-day conditions as described previously (Kasten 2016). Plants were exposed to 30 ppm NO_2_ for 1 h or 20 min. Please refer to (Mayer et al., 2018) and (Kasten et al., 2017) for more details on the fumigation system used. *pDex::AvrRpm1-HA* plants were sprayed until run-off with 30 µM dexamethasone (Sigma) dissolved in 0.1% methanol/0.01% Tween 20. As a negative control, plants were treated with methanol/Tween. Furthermore, we triggered HR-PCD by infection with *P*.*s*.*t*. carrying the avirulence gene *AvrRpm1* as described earlier (Gaupels et al., 2011).

### Fluorescence imaging

The imaging of BG autofluorescence in whole plants and leaves was done as reported before (Mayer et al., 2018). For microscopic examinations the upper side of the leaf was opened by shallow vertical cuts. Subsequently, strips of the adaxial epidermis attached to mesophyll cells were peeled off using forceps and immediately mounted on a microscope slide in water. Occasionally, water was replaced by 50% glycerol solution to induce protoplast shrinking. An Olympus BX61 fluorescence microscope equipped with a XC50 camera served to define the tissue localization of the red and BG fluorescence. Fluorescence was excited with the 365 nm band of the UV laser and detected after passing a UV longpass filter. Additionally, we investigated epidermal peels with a Zeiss LSM 510 Meta CLSM. Here, the microscope setup included a 364 nm excitation laser combined with a 385 nm longpass filter for the detection of BG fluorescence, or a 633 nm excitation laser combined with a 650 nm longpass filter for the detection of red fluorescence. The Zeiss ZEN lite and arivis Vision4D softwares facilitated the preparation of stacks, other figures, and 3D videos.

### 12-Tesla FT-ICR-MS and MS/MS fragmentation analysis

Metabolite extraction and FT-ICR-MS measurements followed published protocols (Kanawati et al., 2017; Mayer et al., 2018). However, for statistical analysis we used in the R software package the beta-binomial test instead of the Wilcoxon rank sum test. MS/MS fragmentation experiments were performed by the use of Argon as an inert collision gas in a collision cell (hexapole) having a relatively higher pressure of 3×10^−6^ mbar, when compared to the base pressure inside the ICR cell (1.5 × 10^−10^ mbar). No nozzle-skimmer dissociation or declustering potential inside the electrospray source was applied. Only slight standard linear ion acceleration of 3 eV was applied (prior to adding over it the MS/MS collision energy for MS/MS). This is necessary to forward the externally generated ions along the different linear ion beam guides toward the ICR cell.

### Mass difference network analysis

Mass difference network (MDN) analysis was performed based on previous work (Tziotis et al., 2011; Moritz et al., 2016; Moritz et al., 2019) by utilizing a home-made algorithm. First, it was determined whether accurate masses were linked by mass difference units corresponding to binding of dehydrogenated sinapic acid (C_11_H_10_O_5_, 222.052823), sinapoylmalate (C_15_H_14_O_9_, 338.063782), 5-hydroxyferulic acid (C_10_H_8_O_5_, 208.037173), 5-hydroxyferuloylmalate (C_14_H_12_O_9_, 324.048132), ferulic acid (C_10_H_8_O_4_, 192.042259) and feruloylmalate (C_14_H_12_O_8_, 308.053217) (Supplemental File S1). Subsequently, MDNs were visualized by the Gephi software (version 0.9.2, https://gephi.org/).

### Laccase assay

50 mM sodium acetate (pH 5.0) containing or not (control) laccase from *Trametes versicolor* (Sigma) at a total activity of 0.004 U was prepared. Subsequently, 50 µl of 50 mM sinapate in DMSO was added to 1 ml laccase solution and the reaction was placed in a shaker incubator at 50 °C for 10 min. After centrifugation for 10 min at 16.000 g and RT 100 µl sample was transferred to a 96-well plate. In a platereader photometer the fluorescence was recorded between 400-600 nm upon excitation at a wavelength of 320 nm.

### UPLC-MS/MS

The phenolic cell wall esters were extracted as reported earlier (Schnitzler et al., 1996) with minor modifications. We weighed 100-150 mg of frozen leaf material in 1.5 mL polypropylene tube and added 11-12 glass pearls. Leaf material was homogenized using the Silamat S6 bead mill (Ivoclar Vivadent) (Mayer et al., 2018). The weight of the tubes containing the glass beads were determined before adding leaf material and after air-drying of the washed insoluble leaf material to assess the amount of crude cell wall in each sample.

For metabolomics we used Ultra Performance Liquid Chromatography (UPLC) Ultra-High Resolution (UHR) tandem quadrupole/Time-of-Flight (QqToF) mass spectrometry (MS). The LC-MS system was composed of an Ultimate 3000RS (Thermo Fisher) coupled to a Bruker Impact II with Apollo II electrospray ionization source (Bruker Daltonic). For the separation of polar cell wall esters we used the hydrophilic interaction liquid chromatography (HILIC) column ACQUITY BEH Amide (Waters, 100 × 2.1 mm i.d. with 1.7 µm particles) with a gradient of solvent A (formic acid/water 99.9 : 0.1 (v/v)) and solvent B (acetonitrile/formic acid 99.9 : 0.1 (v/v)) following the gradient program depicted in Supplemental Table S1.

Mass calibration was performed with the calibration mixture of 50 mL of water, 50 mL 2-propanol, 1 mL NaOH, and 200 µL of formic acid. The sample material was analysed using MS in negative electrospray ionization mode using the following parameters: Nebulizer pressure, 2.0 bar; dry gas flow, 8.0 L/min; dry gas temperature, 200°C; capillary voltage, 3500 V; endplate offset, 500 V; mass range, 20–2000 *m/z*. Smart exclusion feature together with the auto MS/MS acquisition mode was used for the precursor selection. The rolling average is used as a criterion to determine if the precursor is taken for MS/MS. When the difference between the rolling average of the set number of spectra and actual spectra is over the absolute threshold of 500 counts, the precursor is taken for MS/MS. A collision energy from 5 to 20 eV was used to obtain fragmentation of the ions.

The data were processed by the Metaboscape 4.0 software (Bruker). Processed data were given as ”bucket“ list containing the exact compound mass, isotope composition and chemical clusters in each bucket. The detailed parameters for the software are given in the Supplemental Table S1. The specific masses were found from the bucket list and assigned with the sum formula using the build it feature. The condition for the formula assignment was 5.0 mDa tolerance for the precursor mass. The missing values were replaced with the average area value from all samples for the corresponding mass feature (Denkert et al., 2006).

## SUPPLEMENTAL DATA

**Supplemental Figure S1**. Identification of compounds by MS/MS fragmentation analyses.

**Supplemental Figure S2**. Blue and BG fluorescent compounds are localized in protoplasts of the adaxial epidermis.

**Supplemental Table S1**. UPLC-MS/MS settings.

**Supplemental File S1**. Normalized FT-ICR-MS dataset including statistics and outcome of the mass difference calculations.

## ACKNOWLEDGEMENTS

We thank Elisabeth Georgii for support with the statistical analysis. No conflict of interest declared.

## AUTHOR CONTRIBUTIONS

F.G., J.D., J.-P.S., and P.S.-K. designed the experiments. B.K., F.G., M.B., F.H., and I.Z. performed the experiments. B.K., M.B., F.H., and F.M. analyzed the data. F.G., P.S.-K., and D.M. wrote and critically revised the manuscript.

## DATA AVAILABILITY

The data that supports the findings of this study are available in the supplementary material of this article.

## FIGURE LEGENDS

**Movie 1**. Localization of fluorescence in untreated leaf tissues by CLSM. Epidermal peels of air fumigated leaves were analyzed. Blue fluorescence was detected upon UV excitation (excitation at 364 nm/emission at ≥385 nm). Red chlorophyll fluorescence was detected at emission wavelengths ≥650 nm after excitation at 633 nm. Stacks of optical sections were used to prepare 3D videos.

**Movie 2**. Localization of HCD-related BG fluorescence by CLSM. Epidermal peels of NO_2_-fumigated leaves were analyzed. Blue fluorescence was detected upon UV excitation (excitation at 364 nm/emission at ≥385 nm). Red chlorophyll fluorescence was detected at emission wavelengths ≥650 nm after excitation at 633 nm. Stacks of optical sections were used to prepare 3D videos. Note that burst chloroplasts in the mesophyll co-localize with bright fluorescence in collapsed epidermis cells. There is no bright fluorescence visible next to intact chloroplasts.

